# Significantly distinct branches of hierarchical trees: A framework for statistical analysis and applications to biological data

**DOI:** 10.1101/002188

**Authors:** Guoli Sun, Alexander Krasnitz

**Affiliations:** Simons Center for Quantitative Biology, Cold Spring Harbor Laboratory, Cold Spring Harbor, NY 11724, USA; Department of Applied Mathematics and Statistics, Stony Brook University, Stony Brook, NY 11794, USA

## Abstract

**Background:** One of the most common goals of hierarchical clustering is finding those branches of a tree that form quantifiably distinct data subtypes. Achieving this goal in a statistically meaningful way requires (a) a measure of distinctness of a branch and (b) a test to determine the significance of the observed measure, applicable to all branches and across multiple scales of dissimilarity.

**Results:** We formulate a method termed Tree Branches Evaluated Statistically for Tightness (TBEST) for identifying significantly distinct tree branches in hierarchical clusters. For each branch of the tree a measure of distinctness, or tightness, is defined as a rational function of heights, both of the branch and of its parent. A statistical procedure is then developed to determine the significance of the observed values of tightness. We test TBEST as a tool for tree-based data partitioning by applying it to five benchmark datasets, one of them synthetic and the other four each from a different area of biology. For each dataset there is a well-defined partition of the data into classes. In all test cases TBEST performs on par with or better than the existing techniques.

**Conclusions:** Based on our benchmark analysis, TBEST is a tool of choice for detection of significantly distinct branches in hierarchical trees grown from biological data. An R language implementation of the method is available from the Comprehensive R Archive Network: cran.r-project.org/web/packages/TBEST/index.html.

## Background

Hierarchical clustering (HC) is widely used as a method of partitioning data and of identifying meaningful data subsets. Most commonly an application consists of visual examination of the dendrogram and intuitive identification of sub-trees that appear clearly distinct from the rest of the tree. Obviously, results of such qualitative analysis and conclusions from it may be observer-dependent. Quantifying the interpretation of hierarchical trees and introducing mathematically and statistically well-defined criteria for distinctness of sub-trees would therefore be highly beneficial and is the focus of this work.

The need for such quantification was recognized some time ago, and methods have been designed for (a) identifying distinct data subsets while (b) making use of hierarchical tree organization of the data. These methods fall into two categories, depending on whether or not they employ statistical analysis. The simplest approach that does not rely on statistical analysis is a static tree cut, wherein the tree is cut into branches at a given height. This procedure is guaranteed to produce a partition of the data, but provides no way to choose the height at which to cut. Moreover, some partitions cannot be produced by a static cut. Dynamic Tree Cut, or DTC in the following [1], is a more sophisticated recipe, capable of generating partitions not achievable by a static cut. However, DTC partitions depend on the minimal allowed number of leaves in a branch, a user-defined parameter that cannot be determined by the method itself.

In addition, there are methods for choosing a tree partition from considerations of branch distinctness and its statistical significance. Sigclust, or SC in the following [2], is a parametric approach wherein a two-way split of the data is deemed significant if the null hypothesis that the data are drawn from a single multivariate normal distribution is rejected. The method is designed to work in the asymptotic regime, where the dimensionality of the objects being clustered far exceeds the number of the objects. In application to trees SC works in a top-down fashion, by first examining the split at the root node and proceeding from a parent node to its daughter nodes only if the split at the parent node has been found significant. Unlike SC, the sum of the branch lengths method, or SLB in the following [3] is designed specifically for hierarchical trees and utilizes a measure of distinction between two nodes joined at a parent node that is linearly related to the heights of the two daughter nodes and that of the parent. Similarly to SC, SLB adopts a top-down scheme.

A method introduced here is termed Tree Branches Evaluated Statistically for Tightness (TBEST) and shares features with the existing approaches. Like SC and SLB, TBEST employs statistical analysis to identify significantly distinct branches of a hierarchical tree. Similarly to DTC and SLB, it uses tree node heights to assess the distinctness of a tree branch. As with the other three methods, partitions generated by TBEST are not necessarily accessible by a static cut.

At the same time, TBEST differs from the existing designs in several aspects, two of which are critical. First, unlike DTC, SC and SLB, it examines all the tree nodes simultaneously for distinctness. Secondly, unlike SLB, it combines node heights non-linearly to construct a statistic for distinctness that is better able to handle a tree in which distinct branches of approximately equal numbers of leaves occur at different heights. The key properties of all four methods are summarized in Table 1.

**Table 1.** Properties of TBEST and of the three published methods.

| Method | Order of examining the tree | Significance estimated |
| --- | --- | --- |
| TBEST | all internal nodes in parallel | Yes |
| SC | top down | Yes |
| SLB | top down | Yes |
| DTC | top down and bottom up | No |

In the remainder of this work we formulate TBEST and systematically compare its performance to that of DTC, SC and SLB on a number of benchmark datasets originating from a variety of biological sources. In all cases we find that TBEST performs as well as or better than the three published methods. We conclude by discussing generalizations of TBEST and its relation to other aspects of cluster analysis.

## Methods

Consider a set of objects with pair-wise relations given by a dissimilarity matrix. Given a linkage rule, a hierarchical tree can be grown for the set. We will only consider inversion-free linkage rules here. The tree is specified, in addition to its branching structure, by the heights of its nodes. The height of the node quantifies the dissimilarity within the data subset defined by the node. We wish to construct, for each node of the tree, a measure of how distinct the data subset corresponding to the node is from the data set. The special case of the objects being points in a Euclidean space, with the dissimilarities defined as distances between the points, may be used for guidance in this construction. The node height then quantifies the linear extent of the data subset defined by the node. Accordingly, it has been proposed [3] to make the measure of distinctness of a node *n* linear in the difference in heights between a parent *P*(*n*) of n and that of *n* itself. An example of a one-dimensional dataset, tabulated in Additional File 1 and shown in Figure 1, illustrates a difficulty with such construction. Both the subsets shown in blue and in green are clearly distinct from the rest of the data, but the difference in heights between the blue node and its parent is not as great as that between the green node and its parent. Thus, based on the parent to child difference in heights, one would conclude, counter-intuitively, that the blue subset is not nearly as distinct as the green subset. A measure in better agreement with intuition is the relative difference of heights:

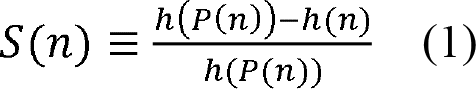

where *h*(*n*) is the height of node *n*. In the following we refer to *S*(*n*) as the tightness of node *n*. In the absence of inversions, the tightness of any node is a number between 0 and 1. In particular, *S*(*n*) = 1 identically if *n* is a leaf. The two subsets highlighted in Figure 1 are nearly equally tight by this measure, despite the disparity in their heights.

**Figure 1.**
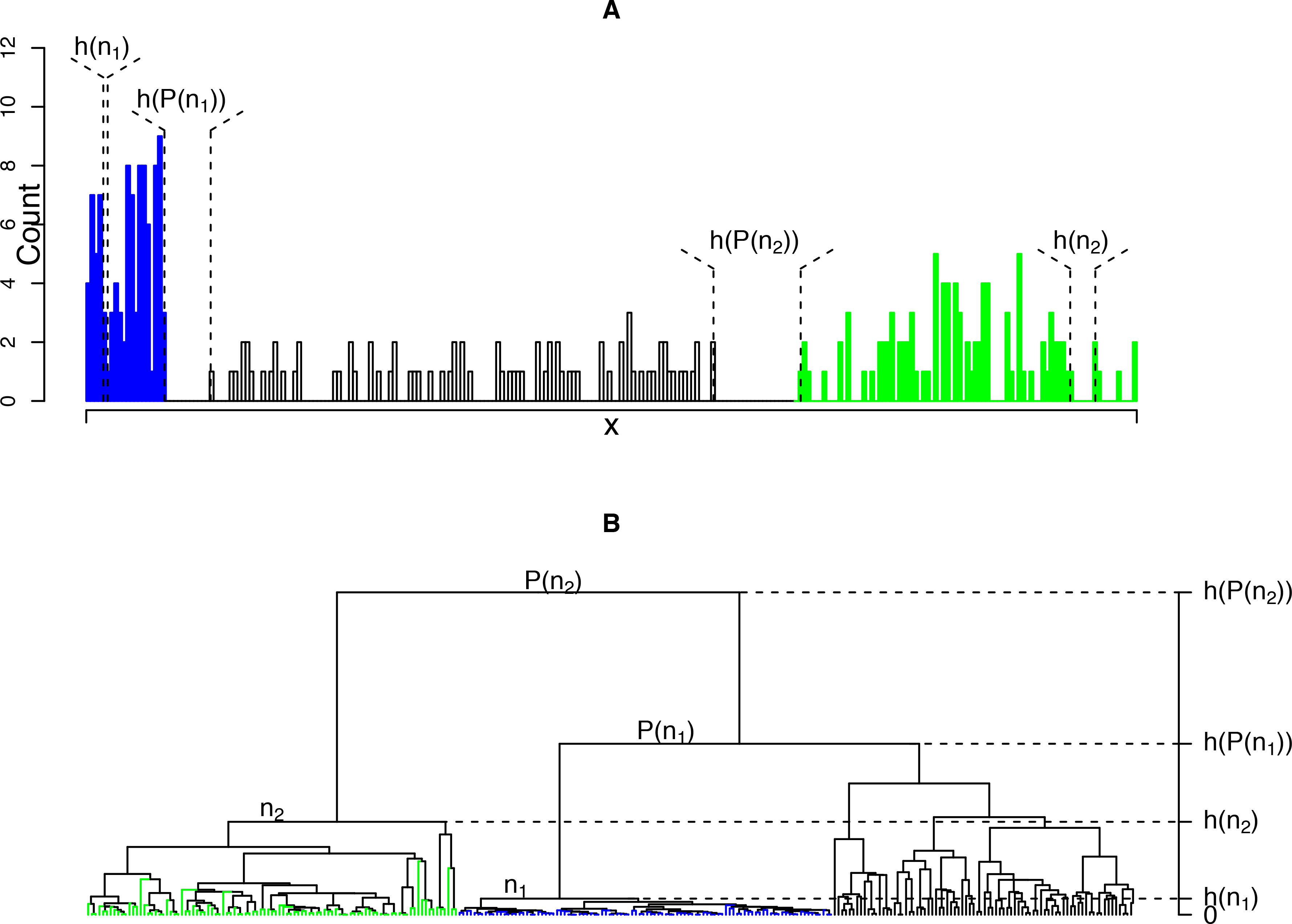
Illustration of the definition of tightness. The data consist of 280 points in one dimension, drawn from a normal mixture with the components *N*(0.5,0.4^2^) (blue), *N*(11,1^2^) (green) and *N*(5,2^2^); (black). **A)** A histogram of the input data. **B)** A hierarchical tree of the input data, grown using the absolute difference of the data values as the dissimilarity measure, and single linkage. Thus, the node heights shown in (**B**) are equal to the corresponding gaps in the data, as indicated in (**A**). Nodes *n*_1_ and *n*_2_ are approximately equally tight.

To enable statistical analysis of tightness, a null distribution of *S*(*n*) is required, for making comparisons with the observed *S*(*n*). This null distribution is obtained by randomizing the dataset from which trees are grown. How such randomization is to be performed depends on the type of the data and on the broader context of the study and cannot be specified in general. For example, if the data matrix represents gene expression, with genes as rows and observations as columns, it may be appropriate to randomize the data by permuting values independently within each row. However, in other situations a more restrictive randomization should be adopted. For example, the elements of a binary data matrix may represent the mutation status at a set of genomic positions (rows) in a collection of genomes (columns). The investigator may wish to randomize the data while preserving both the site mutation frequencies (row sums) and the overall mutation burden within each genome (column sums).

Here we design a general procedure for constructing the null distribution of tightness for any given data randomization scheme. To guide this design, we generated distributions of tightness in trees grown from randomized data for multiple combinations of datasets, definitions of dissimilarity, linkage rules and randomization methods, as listed in Table 2. As Figure 2 and Additional file 2: Figure S1 illustrate, the shapes of these distributions generally depend on the number of leaves and, in most cases examined, the peak of the distribution occurs at higher tightness for smaller number of leaves. The identity *S*(*n*) = 1 for single-leaf nodes is consistent with this observation. We therefore conclude that, for a given observed value of tightness, the appropriate null distribution should be sampled by repeated randomization of the data, growing a tree for each randomization, selecting among its nodes the ones with the numbers of leaves matching the observation, and determining the tightness of these nodes. However, it is not guaranteed that, in any tree grown from randomized data, there will be a unique node with a number of leaves exactly equal to that of the observed node. To resolve this difficulty conservatively, we adopt the following procedure. If, for a given data randomization, the tree contains nodes with the number of leaves exactly as observed, the highest *S*(*n*) computed for these nodes is added to the sample. Otherwise we consider all the nodes with the number of leaves nearest the observed one from above and all those with the number of leaves nearest the observed one from below, and add to the sample the highest *S*(*n*) of any of these nodes.

**Figure 2.**
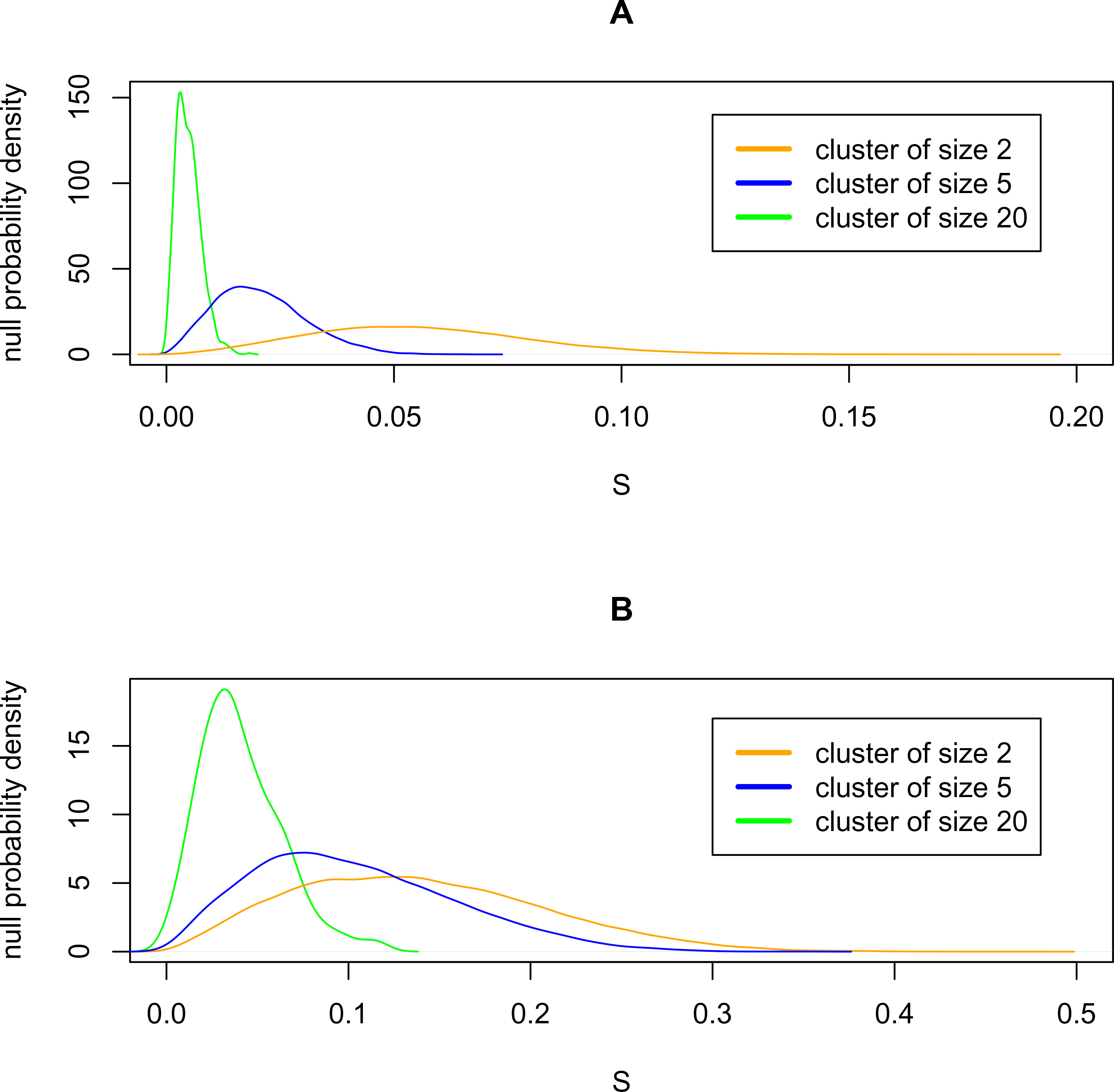
Null distribution of tightness. The null distribution of node tightness *S* depends on the number of leaves. The empirical probability density distributions for the Simulated6 set with (1 - Pearson correlation) dissimilarity – average linkage combination (**A**) and for the Organelles set with (1 - Pearson correlation) dissimilarity – Ward linkage combination (**B**) are shown, for three different values of the number of leaves in each case. Each plot is based on 5000 randomizations of the respective data set.

**Table 2.** Combinations of datasets, dissimilarity, linkage and randomization methods used for testing TBEST.

| Dataset | Dissimilarity | Linkage | Data permutation Method |
| --- | --- | --- | --- |
| Simulated6 | Euclidean | complete | Independently for each coordinate (column) |
|  | (1 - Pearson correlation) | average |  |
| Leukemia | Euclidean | Ward | Independently for each gene (column) |
|  | (1 - Pearson correlation) | average |  |
| T10 | Euclidean | Ward | Independently for each chromosome; identically for all cores (columns) in a chromosome |
|  | (1 - Pearson correlation) | average |  |
| Organelles | (1 - Pearson correlation) | Ward | Independently for each protein (column) |
|  | (1 - Pearson correlation) | average |  |
| Chondrosarcoma | (1 - Spearman correlation) | Ward | Independently for each surface marker (column) |
|  | (1 - Kendall correlation) | average |  |
|  | Manhattan | Ward |  |

With the sampling procedure specified, tests for statistical significance of tightness can be conducted for all the internal nodes of the observed tree, excluding the root, since the latter has no parent. The number of tests is therefore two less than the number of leaves. Due to this multiplicity of tests, higher levels of significance are required for rejection of the null hypotheses for trees with larger numbers of leaves. A straightforward way to handle this requirement would be to increase the size of the sample from the null distribution by performing more randomizations. However, for trees with large numbers of leaves this simple-minded approach may be rendered impractical by computational cost. Instead, higher levels of significance may be accessed by using the extreme-value theory (EVT) to approximate the tail of the null distribution, thereby permitting considerable economy of computational effort [4]. We have used the EVT-based method alongside the more costly purely empirical computation of significance in our benchmark studies reported in the following, and found the two approaches to be in good agreement, as shown in Additional file 2: Figure S2. The *p*-values displayed in the following were computed by applying a multiple-hypotheses correction of the form *p* = 1 − (1 − *p_e_*)*^N^*^-2^, where *p_e_* is the empirical *p*-value and *N* is the number of leaves.

## Results

We evaluated the performance of TBEST in comparison to three published methods of identifying distinct subsets of observations, namely, DTC, SC and SLB. Of the five datasets used in the evaluation one is synthetic, generated to simulate a set of gene expression profiles. The remaining four datasets share two common features: they originate in biological experiments and in each case there is an independently known, biologically meaningful partition of observations into types. We call this known partition “truth”, and the corresponding types the true types, henceforth. The essential properties of the benchmark datasets are summarized in Table 3.

**Table 3.**
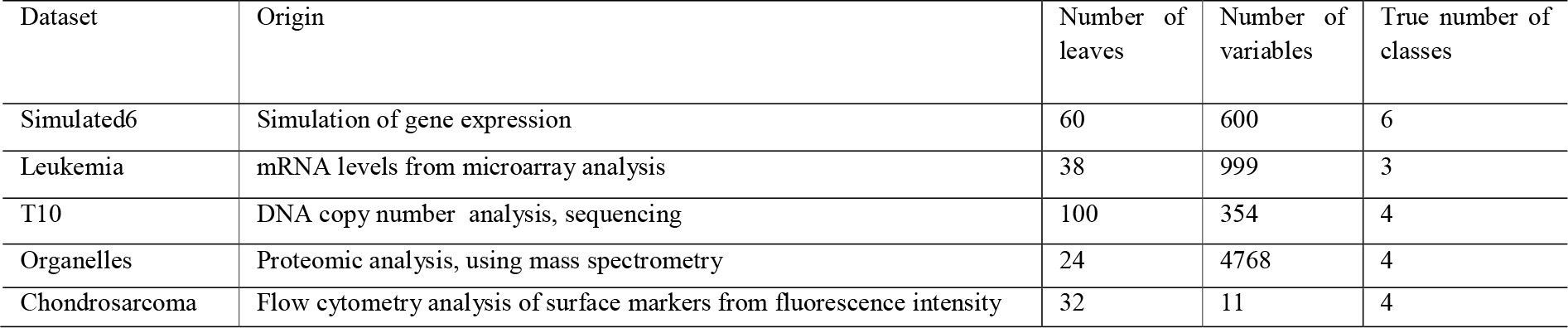
Properties of the five benchmark datasets.

To better judge the performance of TBEST in comparison to the other three algorithms, we considered, for each dataset, more than one combination of dissimilarity and linkage methods used for hierarchical clustering. With the exception of the third benchmark case, randomization of the input data, as required for both TBEST and SLB, consisted of randomly permuting the observed values, independently for each variable. The degree of agreement between a computed partition of the data and the truth is quantified in terms of corrected-for-chance Rand index, or cRI in the following [5]. It should be noted that the subsets of the data identified as distinct by TBEST and the other three techniques by necessity correspond each to a branch of a tree. This, however, is not necessarily the case for the true types, some of which do not correspond to a single branch. As a result, a perfect match between any computed partition and the truth may not be possible, and the maximal attainable value of cRI may be below 1. For this reason, to evaluate the performance of TBEST and the published methods across benchmark datasets, we also identify, for each tree considered, a partition into branches that best matches the truth and determine cRI between that partition and the computed partitions for each of the methods.

### Simulated6

The data are a sample of size 60 in 600 dimensions [6]. The true partition of the data is into six subtypes, with the sizes of 8, 12, 10, 15, 5, and 10. Each of the 600 variables represents a simulation of a gene expression. For 300 of these genes the values are sampled from the same distribution for all subtypes. The remaining 300 genes fall into six non-overlapping subsets of equal size. Each subset corresponds to exactly one subtype, and for that subtype only the genes in the subset are sampled from a distribution that differs from the background.

The comparison between the four algorithms is displayed graphically in Figure 3. For both combinations of dissimilarity and linkage only TBEST and DTC match the truth exactly, while the other two methods either fail to partition the set or do so incompletely. We note that the Euclidean dissimilarity – complete linkage combination results in a particularly challenging tree, which cannot be partitioned correctly by a static cut.

**Figure 3.**
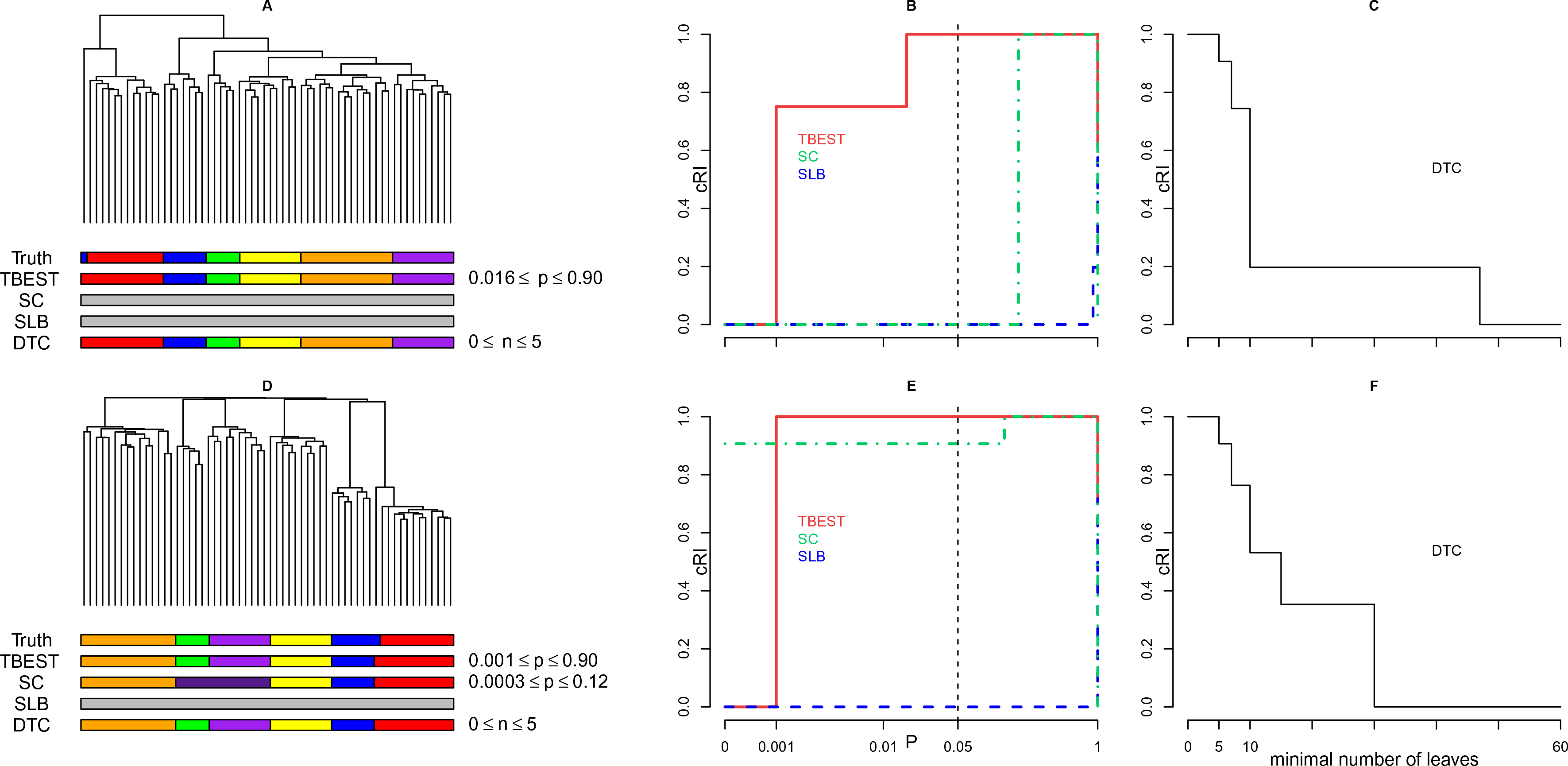
TBEST compared to published methods for Simulated6. Performance comparison of TBEST and the three published methods in Simulated6 dataset for the Euclidean dissimilarity – complete linkage combination (top) and for the (1 - Pearson correlation) dissimilarity – average linkage (bottom). For each combination the left portion (**A** or **D**) shows the corresponding dendrogram, under which then true partition and the partition best matching the truth for each of the methods are shown as color bars. In the middle portion (**B** or **E**), the relative cRI of the computed partition is plotted against the required level of significance *p* for each of the significance-based methods. The customary *p* = 0.05 threshold of significance is shown by a dashed vertical. In the right portion (**C** or **F**), the relative cRI of the computed partition is plotted against the minimal allowed number of leaves for DTC.

### Leukemia

The original Leukemia dataset [7] contained mRNA level values for 6817 genes; this number was reduced to 999 by feature selection [6]. The truth is a partition of patient cases into those of acute myeloid leukemia (AML, 11 cases) and of acute lymphoblastic leukemia (ALL), and a further partition of the ALL subset into the B-cell lineage (19 cases) and the T-cell lineage (8 cases) types. Performance of TBEST is compared with that of the other three methods in Figure 4. For the Ward linkage, two of the significance-based methods, SC and TBEST, attain the highest possible value of the cRI. However, SC only does so with low significance (*p* > 0.33), while TBEST achieves it best performance with high significance (*p* ≈ 2×10^−3^) and maintains performance close to optimal in a wide range of *p*-values. The performance of SLB in this case is similar to that of TBEST, but SLB does not attain the optimum. With the average linkage, TBEST outperforms both SC and SLB throughout the entire range of *p*-values considered and attains optimal performance at high significance. In both cases the performance of DTC is highly sensitive to the minimal allowed size of a branch, especially so for the Ward linkage, where this algorithm attains top performance for sizes between 6 and 10, but performs substantially below the optimum outside this range.

**Figure 4.**
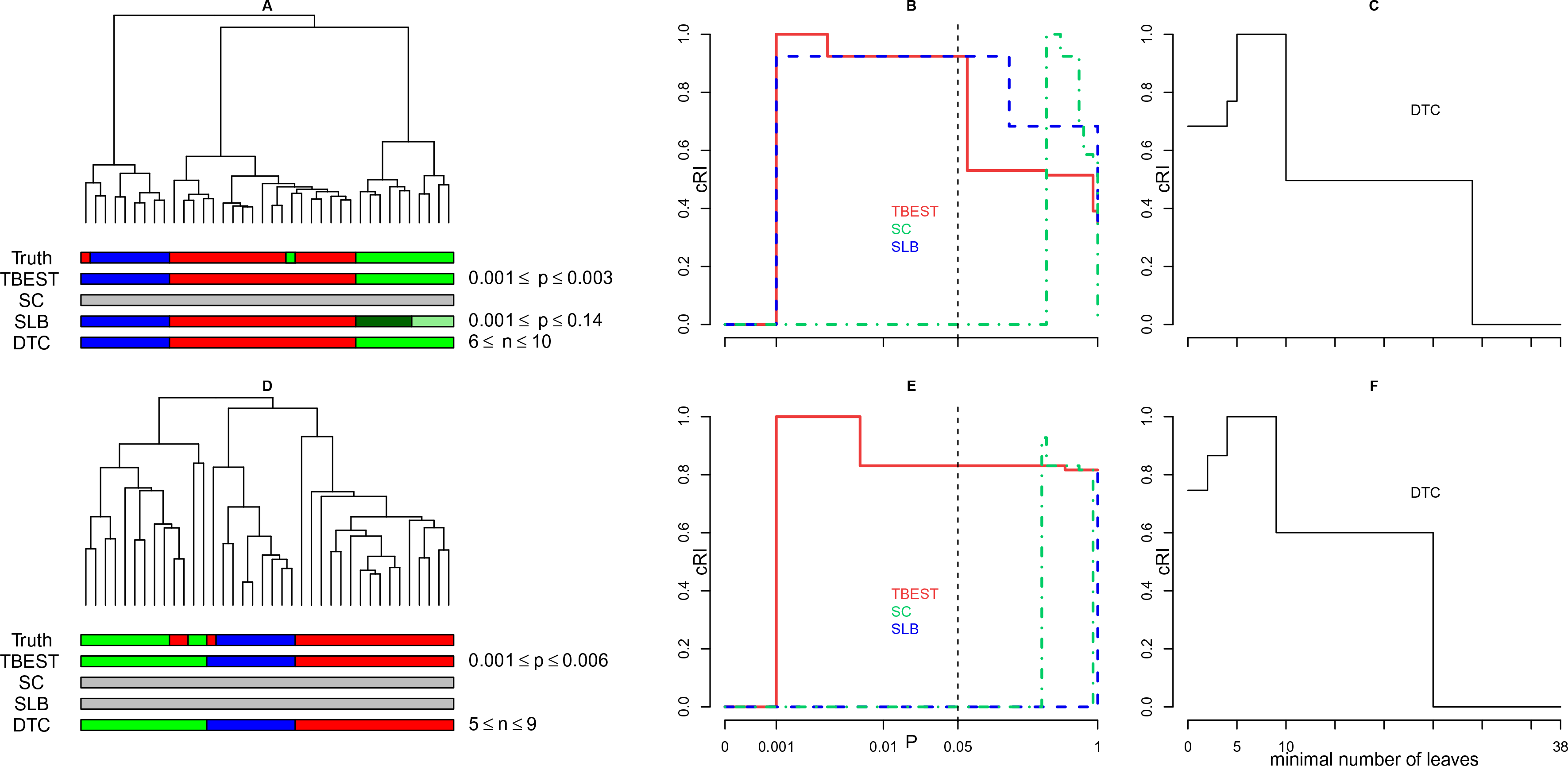
TBEST compared to published methods for Leukemia. Performance comparison of TBEST and the three published methods in Leukemia dataset for the Euclidean dissimilarity – Ward linkage combination (top) and for the (1 - Pearson correlation) dissimilarity – average linkage (bottom). For each combination the left portion (**A** or **D**) shows the corresponding dendrogram, under which then true partition and the partition best matching the truth for each of the methods are shown as color bars. In the middle portion (**B** or **E**), the relative cRI of the computed partition is plotted against the required level of significance *p* for each of the significance-based methods. The customary *p* = 0.05 threshold of significance is shown by a dashed vertical. In the right portion (**C** or **F**), the relative cRI of the computed partition is plotted against the minimal allowed number of leaves for DTC.

### T10

The third benchmark dataset originates from DNA copy number analysis of 100 individual cells harvested from a breast tumor [8]. The true partition in this case is four-way, with the subsets differing from each other by ploidy as determined by cell sorting. The rows of the data matrix correspond each to a cell, the columns correspond each to a pre-defined genomic region of recurrent copy number variation called a core, specified by the sign of variation (gain or loss) and the endpoint positions of the region. The entries in the matrix quantify the extent to which copy number alterations observed in the cells match the cores [9].

There are multiple instances of strong geometric overlap between cores. As a result, the corresponding columns in the data matrix exhibit strong pairwise correlations, positive for cores of equal sign (both gains or both losses), and negative for cores of opposite signs. Consistent with these geometric constraints, the null distribution in this case is generated as follows: the data matrix is divided into sub-matrices by the chromosome number (1,2,…,22,X), and rows are permuted independently within each sub-matrix. The results are illustrated in Figure 5. For the Euclidean dissimilarity - Ward linkage combination only TBEST and SLB identify the true partition, with TBEST succeeding in a broader range of *p*-values. For the (1 – Pearson correlation) dissimilarity - average linkage combination TBEST outperforms the other two significance-based algorithms and matches the truth perfectly in a broad range of *p*-values.

**Figure 5.**
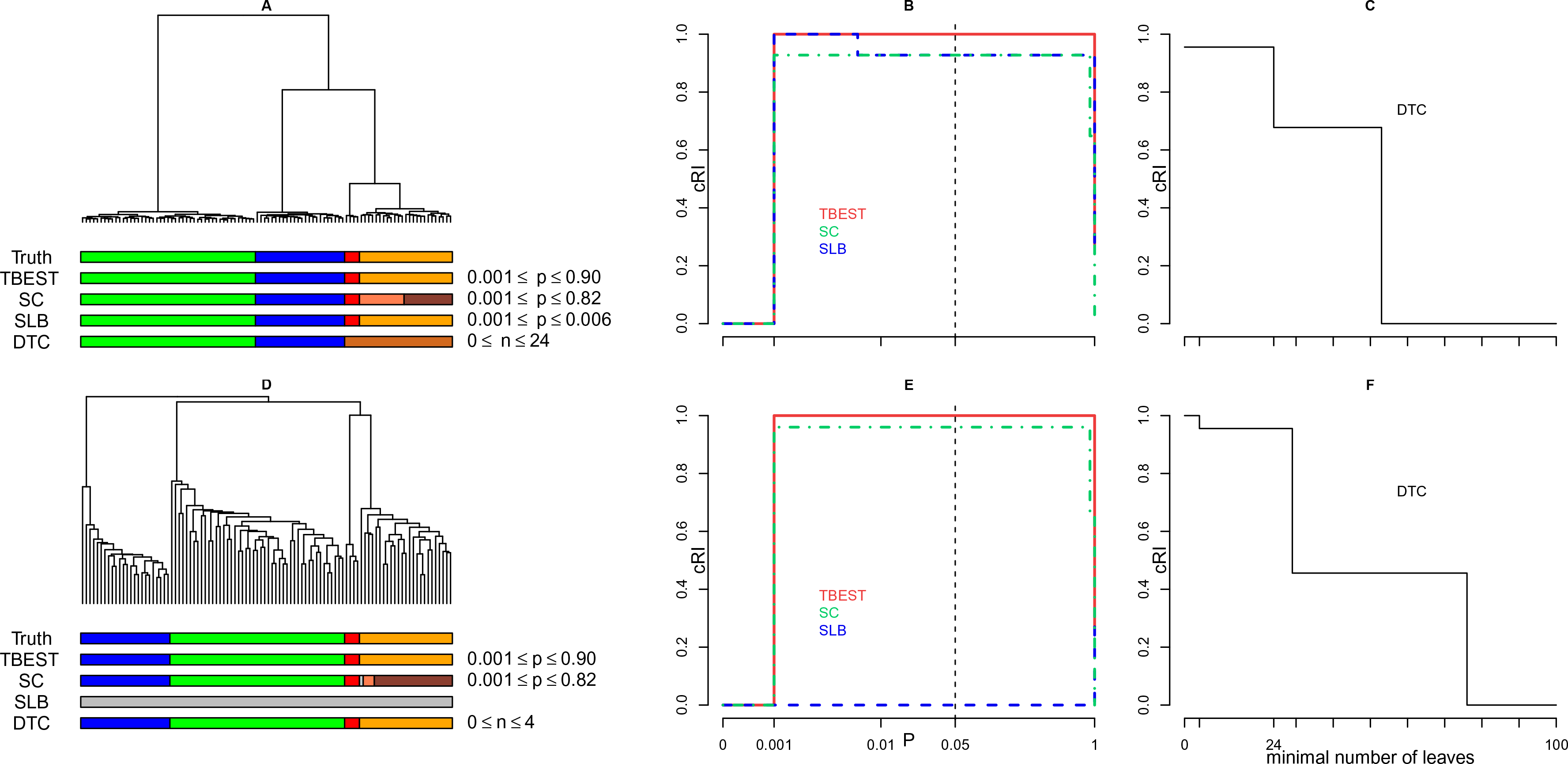
TBEST compared to published methods for T10. Performance comparison of TBEST and the three published methods in T10 dataset for the Euclidean dissimilarity – Ward linkage combination (top) and for the (1 - Pearson correlation) dissimilarity – average linkage (bottom). For each combination the left portion (**A** or **D**) shows the corresponding dendrogram, under which then true partition and the partition best matching the truth for each of the methods are shown as color bars. In the middle portion (**B** or **E**), the relative cRI of the computed partition is plotted against the required level of significance *p* for each of the significance-based methods. The customary *p* = 0.05 threshold of significance is shown by a dashed vertical. In the right portion (**C** or **F**), the relative cRI of the computed partition is plotted against the minimal allowed number of leaves for DTC.

### Organelles

Next, we consider a dataset derived from proteomic analysis of the content of four cellular compartments in each of six mouse tissues. The analysis is based on 4768 protein level readings [10].

The true partition of the data is by the cellular compartment, and the two hierarchical clustering methods considered here both have the branch structure organized by the compartment label, to a good approximation. Of the three significance-based methods compared, only TBEST reproduces the truth to the maximal extent possible for both combinations of dissimilarity and linkage, and it does so stably in the broadest range of the levels of significance (Figure 6).

**Figure 6.**
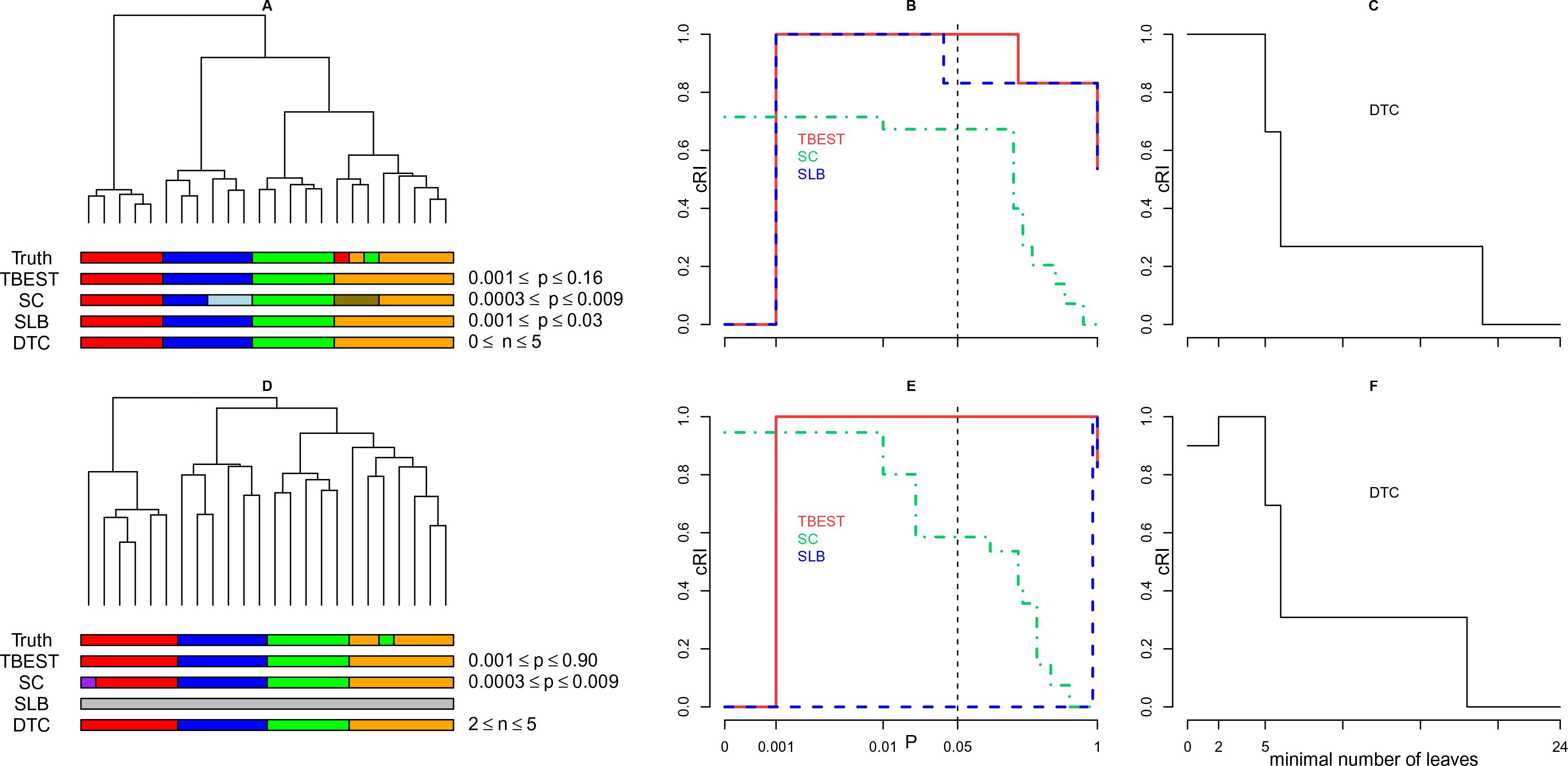
TBEST compared to published methods for Organelles. Performance comparison of TBEST and the three published methods in Organelles dataset for the (1 - Pearson correlation) dissimilarity – Ward linkage combination (top) and for the (1 - Pearson correlation) dissimilarity – average linkage (bottom). For each combination the left portion (**A** or **D**) shows the corresponding dendrogram, under which then true partition and the partition best matching the truth for each of the methods are shown as color bars. In the middle portion (**B** or **E**), the relative cRI of the computed partition is plotted against the required level of significance *p* for each of the significance-based methods. The customary *p* = 0.05 threshold of significance is shown by a dashed vertical. In the right portion (**C** or **F**), the relative cRI of the computed partition is plotted against the minimal allowed number of leaves for DTC.

DTC achieves top performance for the (1 - Pearson correlation) dissimilarity – Ward linkage combination if its minimal allowed number of leaves does not exceed that of the smallest compartment-associated branch of the tree. However, this property is lost for the (1 - Pearson correlation) dissimilarity – average combination where an additional cluster with two leaves is identified by DTC if the minimal number of leaves is set at or below 2.

### Chondrosarcoma

Finally, we discuss the performance of the four methods on a dataset generated by flow cytometry analysis of cells harvested from human tissues and cell lines. Among 34 samples, two samples were identified as multivariate outliers and removed before clustering [11]. The truth is a four-way partition, with three parts corresponding each to a different tissue of origin and the fourth part formed by cells from tumor cell lines.

We have identified three combinations of dissimilarity and linkage for which the tree structure is fully consistent with the true partition and performed comparative analysis for all three, as shown in Figure 7. For two of these combinations ((1 - Spearman correlation) dissimilarity – Ward linkage and (1 - Kendall correlation) dissimilarity – average linkage) partition by TBEST matches the truth in a range of acceptable levels of significance. SLB only does so for the first combination, while SC fails to match the truth. Note the data dimension in this case is 11, and it is smaller than 32, the number of observations. This dataset is therefore outside the range of applicability of SC. For Manhattan dissimilarity – Ward linkage TBEST also matches the truth, albeit at low significance (*p* = 0.1). DTC performs well for the first and third combinations, but only matches the truth in a restricted range of numbers of leaves in the second case.

**Figure 7.**
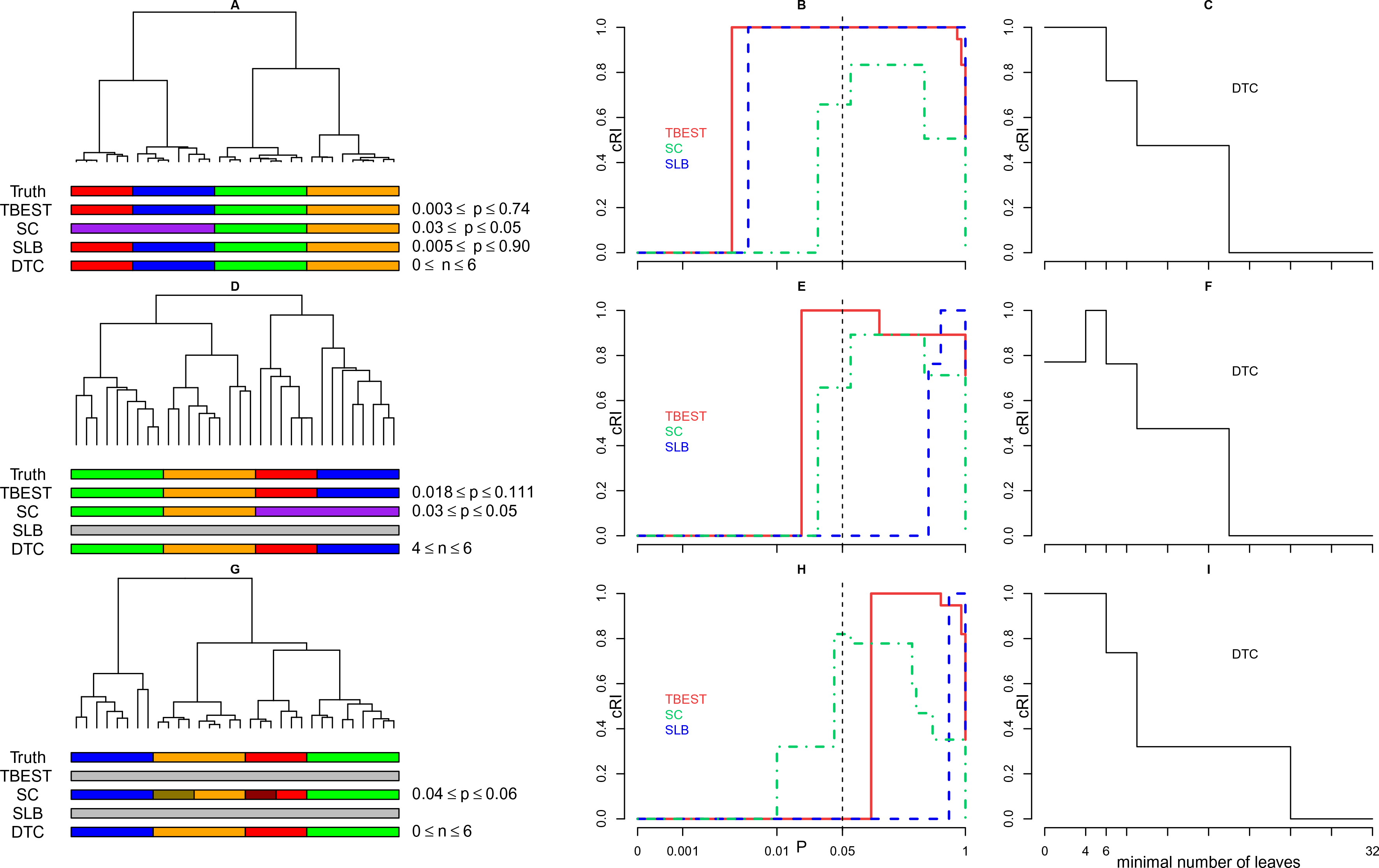
TBEST compared to published methods for Chondrosarcoma. Performance comparison of TBEST and the three published methods in Chondrosarcoma dataset for the (1 - Spearman correlation) dissimilarity – Ward linkage combination (top), (1 - Kendall correlation) dissimilarity – average linkage combination (middle), and Manhattan dissimilarity – Ward linkage (bottom). For each combination the left portion (**A**, **D** or **G**) shows the corresponding dendrogram, under which then true partition and the partition best matching the truth for each of the methods are shown as color bars. In the middle portion (**B**, **E** or **H**), the relative cRI of the computed partition is plotted against the required level of significance *p* for each of the significance-based methods. The customary *p* = 0.05 threshold of significance is shown by a dashed vertical. In the right portion (**C**, **F** or **I**), the relative cRI of the computed partition is plotted against the minimal allowed number of leaves for DTC.

## Discussion and Conclusions

As our test results demonstrate, the performance of TBEST as a tool for data partitioning is equal or superior to that of similar published methods in a variety of biology-related settings. This is true in particular for datasets with underlying tree-like organization, such sets of genomic profiles of individual cancer cells, of the same type as our second benchmark case above. In a work presently in progress we are applying TBEST systematically to a number of datasets of a similar nature. But TBEST also performs well on datasets with no underlying hierarchical structure, such as Simulated6 or Leukemia above. In total, TBEST was able to recover the true partition of the data on par with or better than the published methods in ten out of eleven test cases considered here. We further note that in all but one cases considered the optimal partition of the data by TBEST also was the most significant nontrivial partition. This was not the case for the other significance-based methods included in the comparison.

TBEST can both be applied and formulated more broadly. The applicability of TBEST is not limited to data partitioning that has been our focus here. TBEST can be used for finding all significantly distinct branches of a hierarchical tree, regardless of whether these form a full partition. Further, alternatives to the test statistic of Equation 1 can easily be devised, For example, for any non-leaf node *n* we can introduce

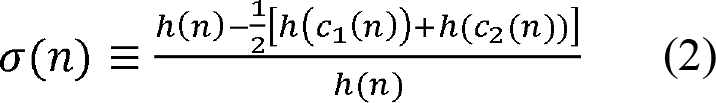

where *c*_1_(*n*), *c*_2_(*n*) are the two children of *n*. While the discussion of these extensions is beyond the scope of this work, an implementation of TBEST as an R language package provides a number of options, both for the definition of tightness and for annotation of significantly distinct branches [12].

Finally, we note that tightness of tree branches is complementary to another important notion in clustering, namely, cluster stability under re-sampling of the input data. The latter property can be analysed in a number of ways, such as bootstrap analysis of trees [13–15] or methods not directly related to trees [6, 16]. Existing work provides examples where both distinctness and stability under resampling are prerequisites of a meaningful partition [17]. Incorporation of TBEST into such combined analysis will be addressed in the future.

## Competing interests

The authors declare that they have no competing interests.

## Authors’ contributions

GS and AK designed the study, wrote software, performed statistical analysis and wrote the manuscript. Both authors read and approved the final manuscript.

## Acknowledgements

We are grateful to M. Wigler for contributing to the early stages of this work and numerous subsequent discussions; to S. Yoon for reading and commenting on the manuscript; to M. Akerman, B. Meunier and J.F. Hoquette for generously sharing their data with us; to K.A. Schlauch for generously providing software.

Funding: This work was supported by the National Institutes of Health grant NIH/1UO1CA168409-01 and by grant 125217 from the Simons Foundation.

---

**Additional file 1 – Dataset displayed in Figure 1**

An Excel file containing a set of 280 positive real values sampled from a mixture of three normal components: *N*(0.5,0.4^2^), *N*(11,1^2^) and *N*(5,2^2^).

**Additional file 2 – Figures S1 and S2.**

A PDF file containing Figure S1, an 11-panel figure illustrating null distribution of tightness and Figure S2, a comparison of empirical *p*-value estimates for tightness to EVT-based estimates.

